# Experimental glaucoma retinal proteomics identifies mutually exclusive and overlapping molecular characteristics with human glaucoma

**DOI:** 10.1101/2020.05.14.095307

**Authors:** Mehdi Mirzaei, Vivek Gupta, Nitin Chitranshi, Liting Deng, Kanishka Pushpitha, Mojdeh Abbasi, Joel Chick, Rashi Rajput, Yunqi Wu, Matthew. J. McKay, Ghasem H Salekdeh, Veer Gupta, Paul A. Haynes, Stuart L. Graham

## Abstract

Current evidence suggests that exposure to chronically induced intraocular pressure (IOP) leads to neurodegenerative changes in the inner retina. This study aimed to determine retinal proteomic alterations in a rat model of glaucoma and compared findings with human retinal proteomics changes in glaucoma reported previously. We developed an experimental glaucoma rat model by subjecting the rats to increased IOP (9.3±0.1 vs 20.8±1.6 mm Hg) by weekly microbead injections into the eye (8 weeks). The retinal tissues were harvested from control and glaucomatous eyes and protein expression changes analysed using multiplexed quantitative proteomics approach. Immunofluorescence was performed for selected protein markers for data validation. Our study identified 4304 proteins in the rat retinas. Out of these, 139 proteins were downregulated (≤0.83) while expression of 109 proteins was upregulated (≥1.2-fold change) under glaucoma conditions (p≤0.05). Computational analysis revealed reduced expression of proteins associated with glutathione metabolism, mitochondrial dysfunction/oxidative phosphorylation, cytoskeleton and actin filament organisation, along with increased expression coagulation cascade, apoptosis, oxidative stress and RNA processing markers. Further functional network analysis highlighted the differential modulation of nuclear receptor signalling, cellular survival, protein synthesis, transport and cellular assembly pathways. Alterations in crystallin family, glutathione metabolism and mitochondrial dysfunction associated proteins shared similarities between the animal model of glaucoma and the human disease condition. In contrast, the activation of the classical complement pathway and upregulation of cholesterol transport proteins, were exclusive to the human glaucoma. These findings provide insights into the neurodegenerative mechanisms that are specifically affected in the retina in response to chronically elevated IOP.

## 1 Introduction

Glaucoma is a common neurodegenerative disease of the retina and is considered as a leading cause of irreversible blindness in the elderly population worldwide. Even though the exact pathophysiology of the disease remains incompletely understood, its association with increased intraocular pressure (IOP) and correlation with the retinal ganglion cell (RGC) death, as well as degeneration of their axons in the optic nerve, is well established [Chitranshi et al., 2018a].

Apart from mechanical or vascular effects of increased IOP, other complex risk factors have increasingly have been associated with the disease onset and progression, including family background, lifestyle, genetics, inflammation, epigenetic factors, secondary neurodegenerative effects of brain disorders, and old age progression [McMonnies, 2017]. Currently, the primary approach to slow down the progression of glaucoma is through reducing the IOP. Nevertheless, thinning of nerve fibre layer and progression of visual field defects continue to be widely reported in the glaucoma patients treated for IOP reduction. This suggests other mechanisms contribute to the disease pathogenesis and progression. There are patients, on the other hand, who exhibit clinical presence of high IOP without corresponding evidence of anatomical or functional damage to the visual system, indicating that elevated IOP may only constitute one of the several variables in the complex pathogenesis of glaucoma [Kass et al., 2002]. To make further advances in glaucoma diagnosis and develop alternative therapeutics, there is a need to systematically define the mechanisms underlying glaucoma. Distinguishing the molecular effects of IOP elevation from a more complex human glaucoma condition, where several other factors might be involved, will help in greater understanding of the disease process.

We recently carried out a comprehensive proteomics study in the post-mortem retinas from human POAG subjects and demonstrated that the pathology is associated with oxidative stress, mitochondrial dysfunction, dysregulation of immune response, disruption of neurotrophic factors and apoptosis activation in the retina [Mirzaei et al., 2017a]. Other proteomics studies in human tissues and animal models have indicated oxidative modification and other post-translational changes in various molecules [Bhattacharya et al., 2006; Tezel et al., 2005]. Enrichment of TNFα signalling pathways and mediation of immune response have also been suggested to play a role in the disease spectrum [Tezel, 2013; Tezel, 2014]. Funke et al. (2016) identified about 600 proteins from the glaucoma and control human retinal tissues using a label free quantitative proteomics approach [Funke et al., 2016]. Proteomics analysis of the non-human primate retinas subjected to experimental glaucoma and optic nerve transection revealed protein changes with little overlap between the two conditions, indicating that mechanisms of RGC damage under these two experimental conditions can be completely different [Stowell et al., 2011]. Andres et al (2017), using the same label-free proteomics technique in an episcleral vein cauterization model, identified 931 proteins that were altered in early phases of IOP increase [Anders et al., 2017]. These studies provided critical insights into the molecular pathogenesis of the disease, however, the latest proteomics technological advances in instrumentation, sample processing and preparation, and bioinformatics analysis tools have enabled us to achieve deeper proteome coverage. In this study, we carried out multiplexed proteomics using TMT SPS-MS3 method and identified about <4300 proteins in the retinas of microbead induced rat model of experimental glaucoma. This is a widely used chronic model of experimental glaucoma and our data reflects that pathways linked to mitochondrial function, cytoskeletal organisation and glutathione metabolism were negatively affected while proteins linked to oxidative stress and apoptotic processes were increased in expression. We analysed this data in the light of previously reported human glaucoma induced changes in the retina and vitreous [Mirzaei et al., 2017a] and this approach permitted us to establish the unique and shared proteins and pathways that are differentially affected under glaucoma conditions.

Identification of these proteins and molecular networks is essential to understand the disease pathophysiology as well as develop target-based drugs and biomarkers for diagnostic applications. This data will serve as an important resource for glaucoma research and will help in comparing the proteomics changes in the retinas of microbead induced chronically increased IOP model with multifactorial human glaucoma condition.

## 2 Results

### 2.1 Quantitative analysis of retinal proteome using multiplexed TMT

A total of 4304 reproducible proteins were identified and quantified from the retinal tissue (1% FDR) - see table 1 (https://data.mendeley.com/research-data/). The quality of the data and reproducibility of the biological replicates across groups were assessed using hierarchical clustering and statistical metrics. We observed great similarity in overall protein abundance distributions of individual biological replicates of both control and glaucoma conditions, confirming the reproducibility of the data for further analysis (Figure 2A). To identify the differentially proteins between glaucoma and control conditions, student’s *t-test* was carried out. Proteins that met the *p*-value cut-off (≤0.05) and at least 20% difference (Table 1) were considered as differentially modulated proteins. This two-step differential analysis of high IOP *vs* control retinas yielded 139 down regulated (p-value ≤ 0.05 and ≤ 0.83-fold change) and 109 up-regulated (p-value ≤ 0.05 and ≥ 1.2-fold change) proteins (Fig 2A, B). A list of top 50 up-and down regulated proteins including their fold changes and their interaction network with p-value cut-off are shown in Fig. 2C, D. Several members of the crystallin family (aa, ba1, ba2, ba4, bb1, bb2, bb3, γs), as well as other proteins including S100a10, S100a4, AHNAK, ANXA1, APOE, CSTS among others were prominently decreased while the expression of coagulation associated proteins including FGG, FGA, FGB, KNG1 and KNG2 were increased in high IOP exposed retinas (Fig. 2C, D).

**Table 1.** The list of proteins identified in the control and glaucoma retinal samples, including differentially expressed proteins identified from pairwise comparison test of Glaucoma vs. Control (student t-test, p-value ≤ 0.05 and ≥ 1.2-fold difference). (https://data.mendeley.com/research-data/)

**Figure 1.**
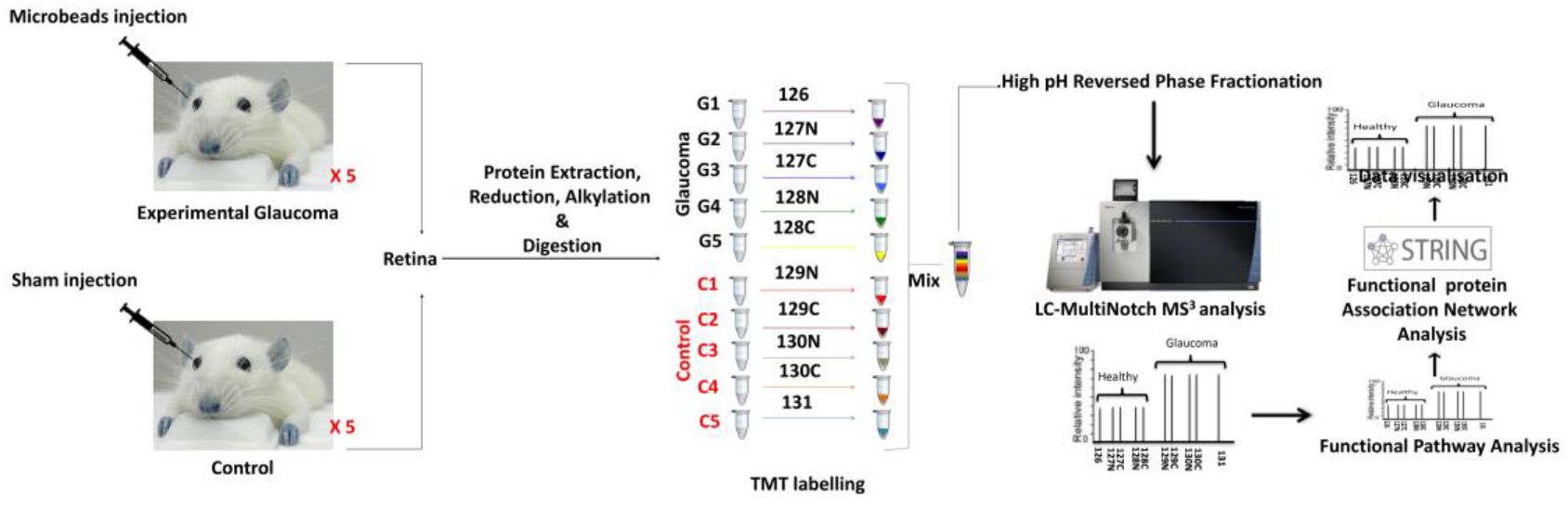
Experimental design and TMT labelling workflow of the experimental glaucoma study. Retinas from control and high IOP rats were harvested (n = 5 each). Extracted proteins from these retinas were alkylated and digested with Lys-C and Trypsin after reduction. The samples were labelled with 10 plex TMT, fractionated and analysed by LC-ESI-MS/MS on ThermoFisher Orbitrap Fusion mass spectrometer (SPS-MS3 method).

**Figure 2.**
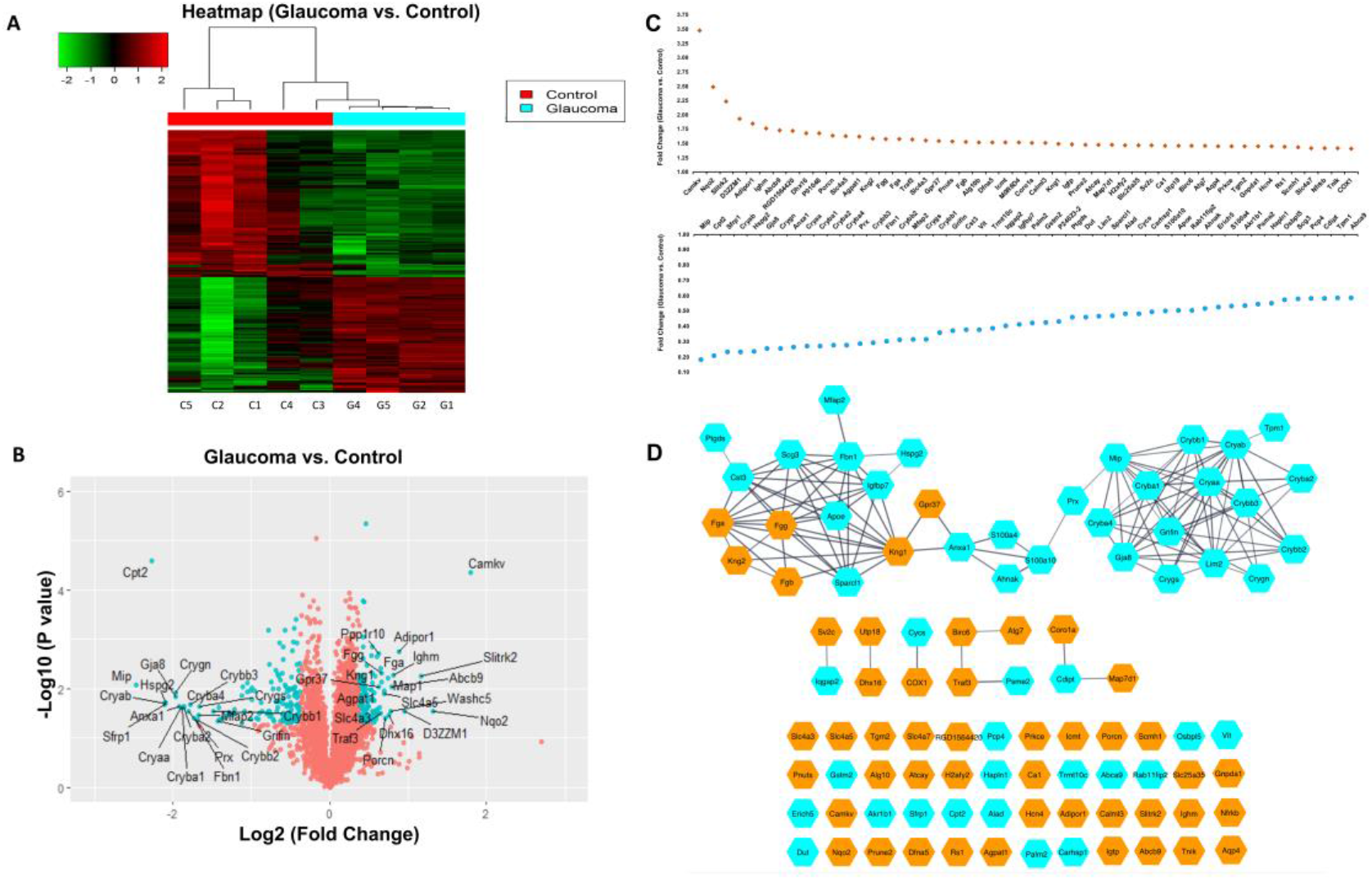
(A) Heatmap (hierarchical clustering) of the log-transformed ratios of differentially expressed proteins from retinal samples (glaucoma vs. control). Red and green colour-coding indicate relative increase or decrease in protein abundance, respectively. (B) Volcano plots representing the expression of differentially regulated proteins. The x-axis represents log2 fold change in abundance in glaucoma versus controls. Proteins above the horizontal dotted line indicate significance ≤ 0.05. Samples that lie in upper and outer quadrants are considered as differentially regulated in glaucomatous conditions. Dots with labelling are the top 50 up and down regulated proteins, respectively. (C) Fold changes of the top 50 up (red colour) and down (green colour) regulated proteins in rat retina with glaucoma are indicated in the graph. (D) Network generated with the top 50 up- and down-regulated proteins using Cytoscape String plugin. The orange and green colours represent the up- and down-regulated proteins respectively.

### 2.2 Differentially regulated cellular pathways and protein networks in glaucomatous retinas

To visualize the protein interaction and networks among these differentially modulated proteins, we generated the retinal protein networks using the STRING and Cytoscape protein-protein interaction analysis tools. From 248 differentially expressed proteins, 21 separate interconnected networks (Fig 3) were identified. Several molecular pathways were reduced in expression, including those involved in actin filament organisation, protein folding, cytoskeleton arrangement, calcium binding, oxidative phosphorylation, and glutathione metabolism. Other networks were increased in expression, such as coagulation, apoptosis and oxidative stress and cell signalling (Fig 3). We performed cellular pathway enrichment and functional protein network analyses on the 248 differentially expressed proteins to understand the molecular mechanisms and biological processes that are altered in the retina in response to exposure to increased IOP. Analysis of the differentially regulated proteins using Ingenuity Pathway Analysis (IPA) revealed that the top enriched disease and implicated biological functions were associated with pathways linked with neurodegenerative processes, modulation of nuclear receptors such as RAR and RXR, Tec kinase and interleukin signalling, mitochondrial dysfunction, fatty acid oxidation and acute phase response signalling (Fig. 4). Overall, the combination of canonical IPA pathway and STRING/ Cytoscape protein-protein network analysis allowed us to visualize the most significantly differentially affected biological processes in the retina of experimental glaucoma model.

**Figure 3.**
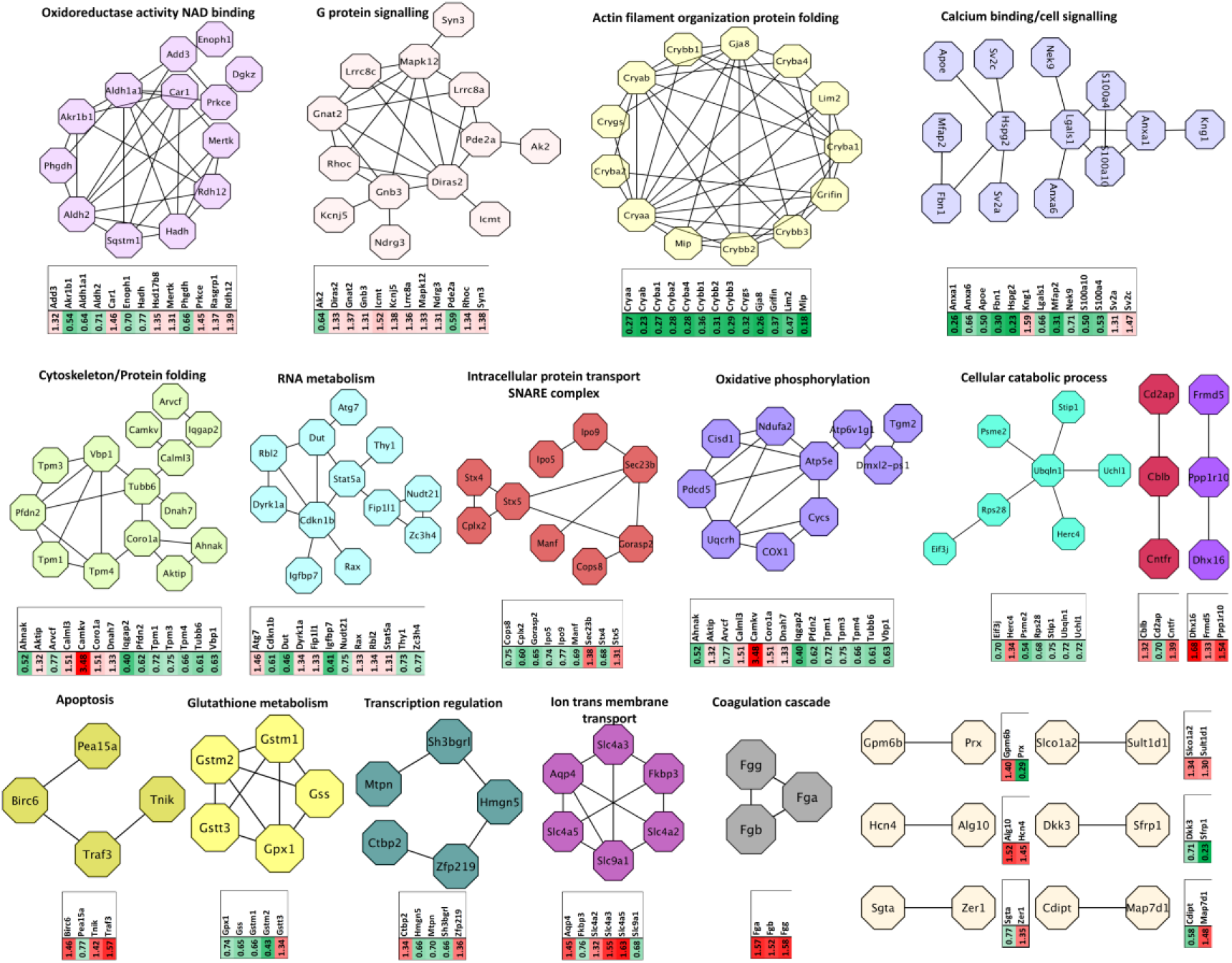
Functional interaction networks analysed by the String Cytoscape plugin. 143 differentially expressed proteins were in generated pathways. Network nodes are labelled with gene symbols, and the corresponding fold changes are indicated below using mini heatmaps. The enriched networks such as oxidoreductase activity, G-protein signalling, cytoskeletal organisation, Ca^2+^ binding, oxidative phosphorylation, RNA metabolism are shown (red-upregulated; Green-downregulated).

**Figure 4.**
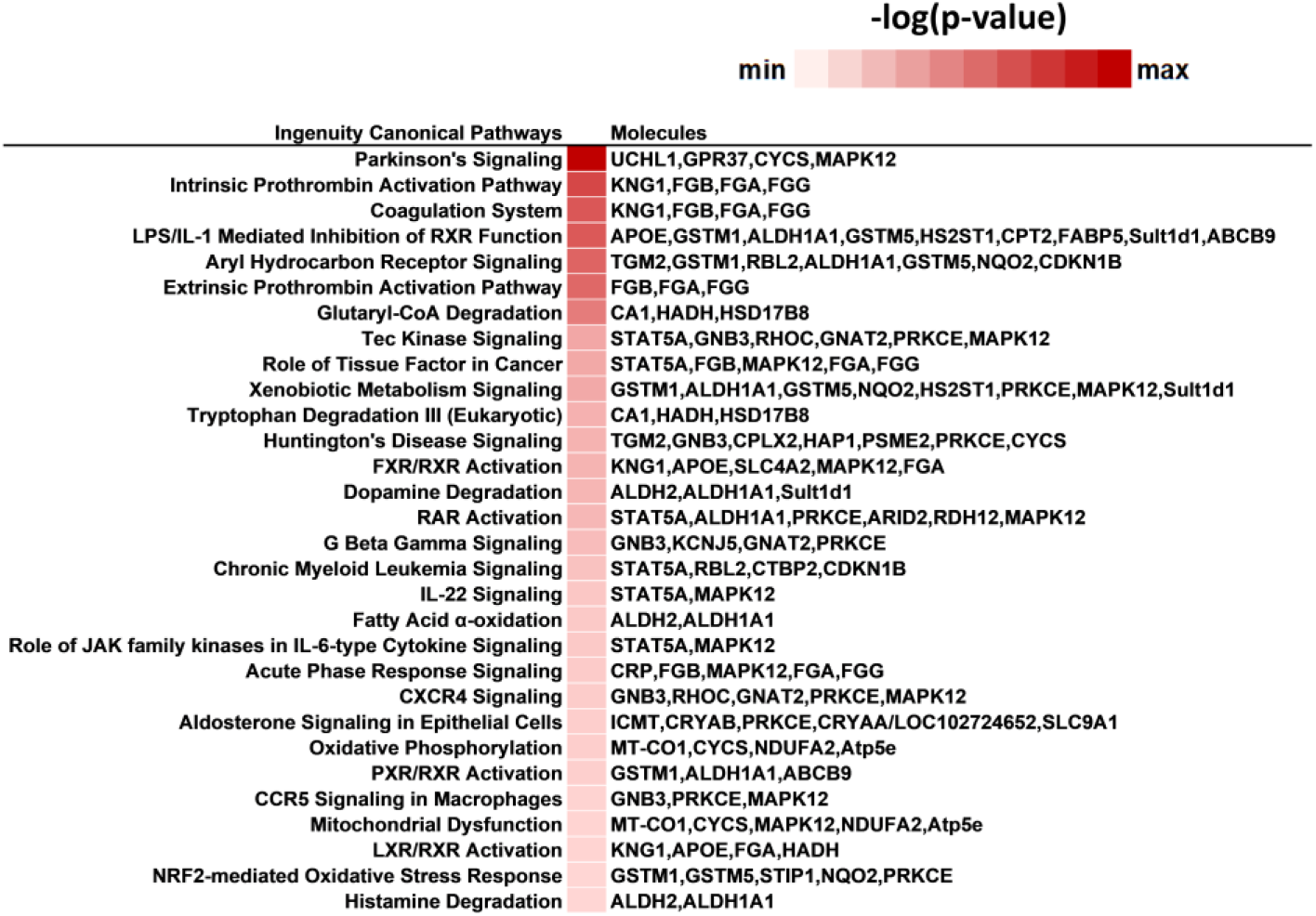
The top 30 canonical pathways enriched from IPA analysis of differentially regulated proteins in retina (glaucoma vs. control). Heatmap of the log transformed abundance ratios representing top disease related biological pathways and differentially affected associated markers in the retina in high IOP exposed eyes. The red colour indicates the significance of the functional enrichment. Proteins in specific pathways are also listed

### 2.3 Redox regulation and oxidative phosphorylation in glaucoma

Mitochondrial dysregulation and impaired oxidative phosphorylation have been implicated strongly in glaucoma pathogenesis [Lee et al., 2011]. A major role of mitochondria is the generation of ATP through oxidative phosphorylation and the regulation of cell death by apoptosis [Hüttemann et al., 2007; Kong et al., 2009]. High energy requirements of the RGCs make them particularly susceptible to anomalies associated with mitochondrial function. We observed reduced abundance of mitochondrial ATP synthase (ATP 5E) and V-type proton ATPase (ATP6v1g1). The oxidative phosphorylation and mitochondrial proteins COX1, Pdcd5, Tgm2 and Ndufa2 were also identified in the list of differentially expressed proteins in glaucoma condition (Fig. 3). It is well documented that IOP increase in glaucoma induces oxidative stress in RGCs through reduced activity of key antioxidant enzymes (e.g; glutathione peroxidases) which are responsible for fighting against oxidative stress [Moreno et al., 2004]. Stress associated proteins such as glutathione metabolic proteins, which are involved in mediating protection of cells against oxidative stress by neutralising highly reactive free radical species, namely GPX1, GSS, GSTM1/2, were significantly downregulated while GSTT3 was observed to be upregulated in glaucoma retinas. Mitochondrial impairment may alter NADPH levels and has been shown to contribute to dysregulated oxidoreductase activity in the cells. In this study we identified that the retinal tissues exposed to high IOP demonstrated reduced levels of aldose reductase (Akr1b1), aldehyde dehydrogenase (Aldh1a1, Aldh2), enolase (Enoph1), acyl dehydrogenase (Hadh) and phosphoglycerate dehydrogenase (Phgdh) enzyme subunits. We also observed upregulation of carbonic anhydrase (Car1), retinal dehydrogenase (RDH12), protein kinase C (Prkce) and NAD dependent 17 β-dehydrogenase (Hsd17b8) enzymes (Fig. 3). These proteins are involved in regulating oxidoreductase activity and NAD binding suggesting that these pathways were negatively impacted in the retinas in response to chronic exposure to high IOP (Fig. 3 and Table 1).

### 2.4 Effects of glaucoma on actin filament organisation and cytoskeletal dynamics

The internal scaffolding of cells is affected in glaucoma with trabecular meshwork, optic nerve and lamina cribrosa exhibiting profound reorganisation of the actin filament networks [Clark et al., 1995; Hoare et al., 2009; Job et al., 2010]. The internal architecture is particularly important to maintain the biomechanical properties and axonal support. We observed consistent downregulation of the proteins associated with actin skeletal organisation and protein folding in the retina under glaucomatous conditions. We identified 9 members of the crystallin family that were downregulated in glaucoma condition (Cryaa, Cryba2, Crygs, Cryab, Crybb1, Cryba4, Cryba1, Crybb2, Crybb3) (Fig. 3, Fig. 5, Fig 6). Crystallins are primarily heat shock proteins that are involved in regulating the actin and intermediate filament cytoskeleton dynamics and protect the cells against stress[Launay et al., 2006]. Reduced levels of Connexin 50 (Gja8) and Griffin cell junction proteins that are involved in mediating cell-cell interactions and cytoskeletal binding were observed in the retinal tissues subjected to high IOP. Our proteomics datasets also included proteins associated with cytoskeleton arrangement and protein folding. Tropomysin isoforms (Tpm1, Tpm3, Tpm 4), which are considered primary regulators of F-actin cytoskeleton, were downregulated in the retinas under glaucomatous conditions. As a matter of interest, pseudokinase Camkv, that may act as scaffold and spatial anchor for cytoskeletal proteins and is also involved in dendrite physiology, was prominently upregulated in the retinas [Jacobsen and Murphy, 2017]. Similarly, Calml3 and Coro1a were overexpressed under the glaucomatous conditions; Calml3 has previously been shown to be differentially regulated in POAG [Liu et al., 2013].

**Figure 5.**
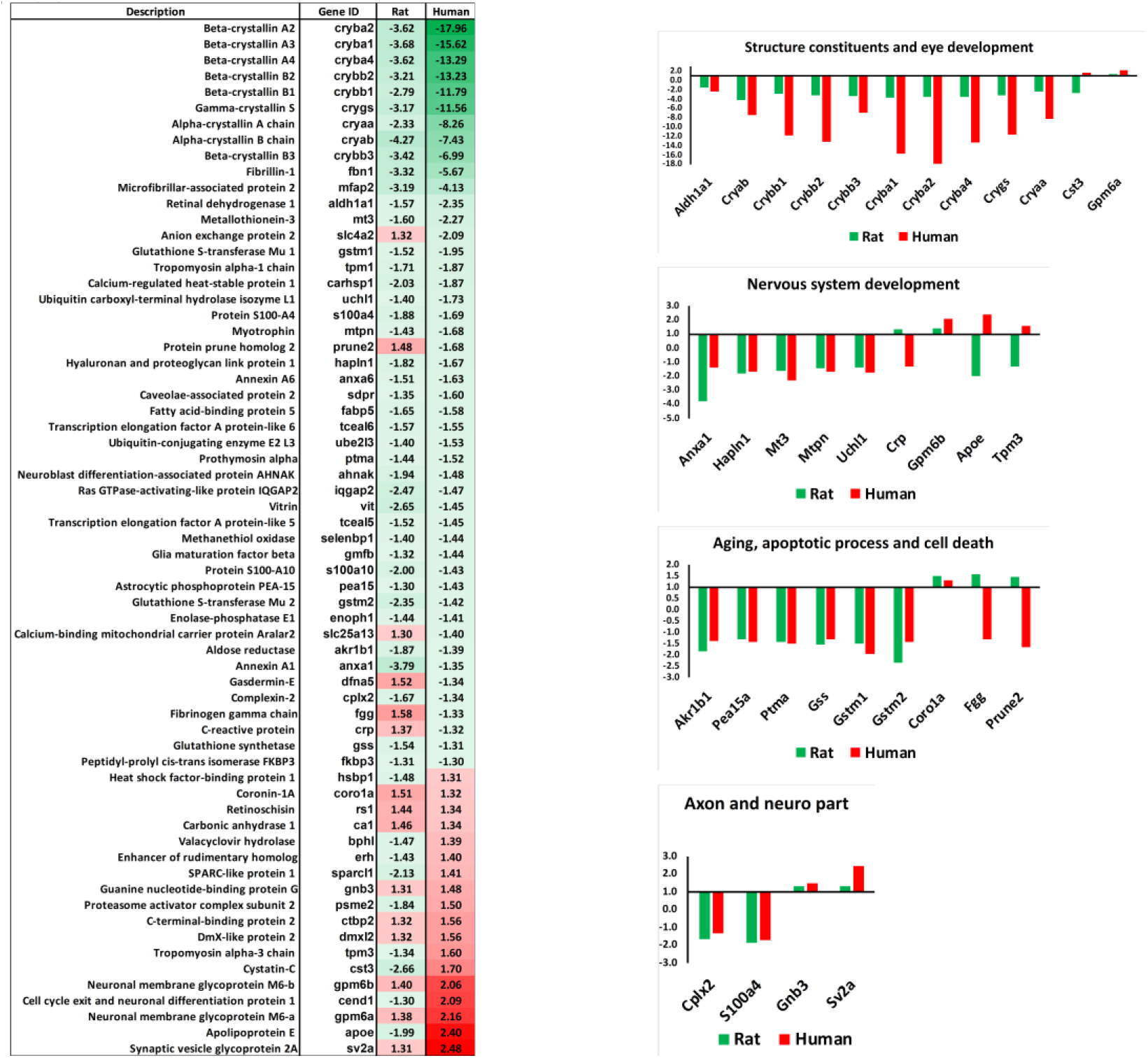
**(A)** Heatmap generated with the common regulated proteins in both rat and human retina with glaucoma. Red and green colours indicate relative increase or decrease (glaucoma vs control) in protein abundance, respectively. **(B)** The bar graphs indicated 4 main pathways in which these common proteins are involved. The regulation patterns of proteins are also revealed.

**Figure 6.**
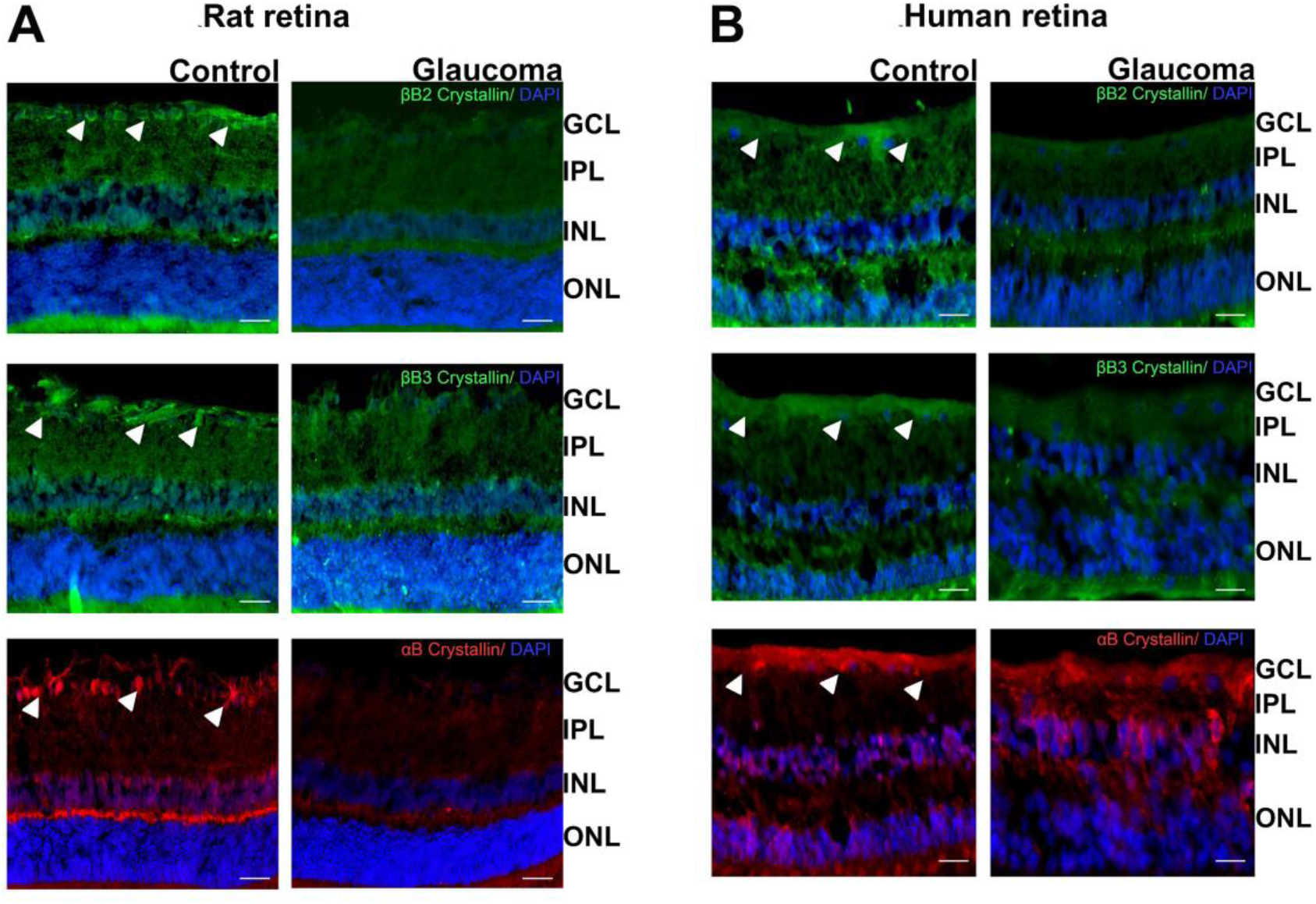
**(A)** Immunostaining of the retinal sections from controlled and glaucoma rats illustrating βB2-Crystallin (Green – Alexa Fluor 488), βB3-Crystallin (Green - FITC) and αB Crystallin (Red – Cy3) expression (Blue-DAPI) (n=3 each). **(B)** Immunostaining of the retinal sections from controlled and glaucoma human postmortem tissues illustrating βB2-Crystallin (Green – Alexa Fluor 488), βB3-Crystallin (Green - FITC) and αB Crystallin (Red – Cy3) expression (Blue-DAPI) (n=3 each). Scale=50μm.

### 2.5 Impaired RNA metabolism and SNARE complex machinery

RNA metabolism was another major cellular process that was impacted in the retinas exposed to glaucomatous injury. We identified six upregulated and six downregulated proteins that have previously been implicated in regulating RNA metabolism, suggesting that RNA processing was negatively impacted. Atg7 has been shown to mediate axonal preservation [Komatsu et al., 2007] and was overrepresented in glaucomatous retinas. Stat5a levels were also enhanced in response to high IOP injury, which corroborates the previous reports of increased levels of Stat5a mRNA in RGCs in response to experimental glaucoma [Wang et al., 2010]. Igfbp7 levels, in contrast, were significantly downregulated under high IOP. This protein has also been shown to be suppressed in response to hypoxia in the retinas in a mouse model of oxygen induced retinopathy [Ishikawa et al., 2010].

In addition to the RNA metabolism, our data suggests significant perturbation of SNARE machinery associated proteins that regulate formation of membrane vesicles as well as the timing and specificity of vesicular fusion [Manca et al., 2019; McKay et al., 2013]. Myocilin has been suggested as a part of the SNARE like complex based on its homology with Q-SNAREs [Dismuke et al., 2012]. We identified 9 proteins associated with the intracellular SNARE transport complex to be differentially regulated in the retinas of high IOP exposed animals. Sec23b and Stx5 proteins which are involved in ER-Golgi transport [Liang et al., 2018] were observed to be upregulated (Fig. 3). On the other hand, Importin proteins (IPO5 and 9), that mediate energy dependent transportation through the nuclear pore, were downregulated [Alqawlaq et al., 2012] along with other SNARE proteins such as Cops8, Cplx2, Gorasp2, Manf and Stx4. Perturbations in SNARE machinery might play a role in dysregulation of intracellular processes including autophagosome fusion with lysosomes, leading to impaired cargo transport and recycling, which are pathological processes implicated in glaucoma [Frost et al., 2014; Porter et al., 2013; Swarup and Sayyad, 2018].

### 2.6 Molecular overlap of protein networks in human and experimental glaucoma

The rat experimental glaucoma data obtained in this study was compared with our previously published [Mirzaei et al., 2017a] retinal and vitreous proteomics data from human POAG tissues. Pathway analysis revealed 65 proteins that were common between the two sets and differentially modulated. A majority of the common differentially expressed proteins were downregulated under both these conditions. Overall, fifty proteins shared a similar expression pattern, amongst these 41 were down- and 9 up-regulated across both systems. Further analysis indicated that within the downregulated set, 9 proteins belonged to the crystallin family that is associated with protein folding (Cryaa, Cryba2, Crygs, Cryab, Crybb1, Cryba4, Cryba1, Crybb2, Crybb3), 3 members to glutathione synthesis pathway (gss, gstm1, gstm2), two were Annexin proteins (anaxa6, anaxa1), and two were S100 proteins (s100a4, s100a10) (Fig. 5A, B). To explore the differential regulation of crystallins, which was identified as one of the most prominent protein groups modulated across rat and human data sets, we subjected both rat and human retinal sections to immunofluorescence analysis using selected crystallin antibodies (αβ, βB2 and βB3) (Fig. 6). The results validated our MS findings and demonstrated noticeable downregulation of expression of each of these isoforms in glaucoma conditions. The downregulation was particularly localised to the GCL in the inner retina, a region that is preferentially affected in glaucoma. The shared up-regulated proteins included glycoprotein m6a (gpm6a), glycoprotein m6b (gpm6b), synaptic vesicle glycoprotein 2A (sv2), DmX-like protein 2 (dmxl2), C-terminal-binding protein 2 (ctbp2), G Protein Subunit β 3 (gnb3), carbonic anhydrase 1 (ca1), retinoschisin (rs1), and coronin1A (coro1a) (Fig. 5A). However, this also means that a substantial number of uniquely affected pathways were also identified in both the human and rat experimental glaucoma conditions. Another noticeable observation in these sets was with respect to the 15 proteins that were differentially modulated but followed an opposite pattern of abundance. Prominent amongst these were fibrinogen subunit (fgg), c-reactive protein (crp), proteasome activator (psme2) and apolipoprotein E, listed in 5A. With this approach, we noted shared differential regulation of proteins implying similar changes in both the data sets in molecular pathways that are involved in mediating retinal structure and development, ageing, apoptotic processes and regulating axonal health. (Fig. 5B).

## 3 Discussion

This study aimed to investigate the molecular basis of glaucoma pathogenesis by taking a systems-level perspective of the experimental glaucoma retinal proteome in microbead model using an unbiased quantitative proteomics approach. We analysed molecular similarities and difference between a glaucoma model and the human POAG condition, by comparing the animal retinal proteome against our previously published human data [Mirzaei et al., 2017a]. To our knowledge, this is the most comprehensive retinal proteome profile in a chronic model of experimental glaucoma, and provides validation of several protein markers recently identified in human glaucoma tissues [Mirzaei et al., 2017a]. Over 4300 proteins were identified with expression of 248 proteins differentially modulated in retina in response to sustained exposure to increased IOP. The study revealed that key pathways involved in cellular stress and tissue homeostasis were affected in the retina under high IOP conditions. Using a combination of functional pathway and protein-protein interaction analysis tools enabled us to identify the specific pathways, which were down regulated or induced in animal model of high IOP. In agreement with human data, this study revealed down regulation of proteins associated with oxidative phosphorylation, glutathione biosynthesis, protein folding, actin filament organisation, and cytoskeleton protein networks. Coagulation cascade was amongst the mutually upregulated pathways in both human and animal model. In accordance with the complex nature of human glaucoma condition, the study revealed that a large series of biochemical pathways were affected in the human glaucoma retina in contrast to the high IOP exposed rat model.

In this study, we identified 11 members of crystallin family as significantly down regulated in glaucoma samples. All these crystallins were embodied in the top 50 down regulated proteins as presented in figure 2C. The STRING protein interaction analysis revealed that crystallins are associated with actin filament organisation and protein folding biological processes. Crystallins belong to the family of small heat shock proteins and consist of three main sub families (α, β, and γ crystallins), which are expressed in RGCs [Liedtke et al., 2007; Piri et al., 2007]. Of these 11 proteins, 6 belonged to the β (CRYBA1, CRYBA2, CRYBA4, CRYBB1, CRYBB3, CRYBB2, CRYBB1), 3 to α (CRYAA, CRYAA-isoform 2 and CRYAB) and 2 to the γ family (CRYGN and CRYGS). Differential expression of crystallins has been reported in number of neurodegenerative disorders including glaucoma [Piri et al., 2007; Prokosch et al., 2013]. Human glaucoma patients have been reported to exhibit increased titer values of antibodies against small HSPs [Tezel et al., 1998]. Crystallins play a role as a molecular scaffold and potential modulator of important signalling molecules in glaucoma, and thus are integral to the process of glaucomatous neurodegeneration [Tezel et al., 1998]. Immunofluorescence analysis of the crystallin expression changes revealed a consistent downregulation of various crystallin isoforms in both rat and human tissues. This downregulation was particularly localised to the GCL region and indicates that crystallin changes in glaucoma are likely induced by exposure to chronically elevated IOP.

In addition to the crystallins, another network associated with cytoskeleton and protein folding that was differentially modulated was the down regulation of 3 tropomysosin members (TPM1, TPM3 and TPM4) along with reduction in protein levels of TUBB6, VBP1, PFDN2, IQGAP2, ARVCF and AHNAK. In contrast, cytoskeletal proteins such as CORO1A and DNAH7, and signalling proteins such as AKTIP, CAlM3 and CAMKV were up regulated. This is consistent with previous studies that have indicated reduction in the levels of Camkv protein in rat model of ischemia reperfusion injury [Tian et al., 2014].

Besides modulation of heat shock and cytoskeletal proteins, Ca^2+^ binding proteins such as S100A10, S100, FBN1 and LGALS1 were identified to be downregulated and these proteins interact with other downregulated proteins such as ANAX6, ANAX1, APOE, NEK9, MFAP2, and HSPG2. This interconnected network was largely downregulated, however, synaptic vesicle glycoproteins 2A and 2C, that have a direct interaction with HSPG2, and KNG1 that interacts with Anxa1, were significantly enriched.

Increased expression of proteins associated with coagulation cascade (FGG, FGB and FGA) was also evident in glaucomatous retinas. Activation of coagulation cascade is associated with inflammatory processes in a number of neurodegenerative diseases, including glaucoma. The mechanisms by which the coagulation system is altered comprises enhanced synthesis and activation of coagulant proteins, decreased synthesis of anticoagulants, and suppression of fibrinolysis. However, not only does inflammation activate coagulation, but this may lead to a vicious cycle, where coagulation in turn perpetuates inflammatory response [Levi and van der Poll, 2010]. A prominent downregulation of oxidative phosphorylation enzymes was detected in the rat retinas, similar to the mitochondrial suppression demarcated by downregulation of electron transport chain and mitochondrial ribosomal proteins reported in human glaucoma tissues [Mirzaei et al., 2017a]. Mitochondrial compromise has been extensively reported in animal models and in human glaucoma condition [Chrysostomou et al., 2013; Munemasa et al., 2010]. RGCs, being metabolically highly active, constitute a cell population that is decidedly susceptible to mitochondrial failure. Modulating mitochondrial integrity, such as by inducing DRP1 inhibition or CoQ10 treatment, has been shown to rescue RGCs in glaucoma [Kim et al., 2015; Lee et al., 2014; Lee et al., 2011]. Mitochondrial dysfunction, in turn, affects the NAD/NADH levels and here we observed downregulation of certain key redox regulatory enzymes in response to high IOP exposure, such as aldehyde dehydrogenase, enolase, aldose reductase, acyl dehydrogenase, and phosphoglycerate dehydrogenase. We have previously reported downregulation of 13 subunits of the NADH dehydrogenase: ubiquinone oxidoreductase complex 1 in the human POAG, and these findings together may reflect a failure of the tissues to maintain redox homeostasis in glaucoma. Williams et al. (2017) demonstrated that administration of the NAD precursor niacin, and gene therapy driving expression of Nmnat1, a key NAD^+^-producing enzyme, modulated mitochondrial vulnerability and prevented experimental glaucoma in ageing animals [Williams et al., 2017].

In a network associated with ion transport across membranes, 3 members of the SLC4 gene family were upregulated in glaucoma; SLC4A2, SLC4A3 and SLC4A5. The modulation of the SLC4 gene family has been linked to ocular and corneal diseases in humans. Mutations in SLC4A4 in addition to their role in renal acidosis are implicated in ocular anomalies including glaucoma [Horita et al., 2018]. Furthermore, 3 proteins associated with apoptotic pathways (BIRC6, TRAF3 and TNIK) were recognized as enriched in glaucoma conditions. BIRC6 protein (baculovirus IAP repeat containing protein 6) is a member of the inhibitor of apoptosis gene family, which encodes negative regulatory proteins that play a preventative role in apoptotic cell death. The proteolytic activity of caspases is tightly controlled by the family of inhibitors of apoptosis which are normally induced in the chronic inflammation in cancers and neurodegenerative diseases. TRAF proteins are essential components of signalling pathways activated by TNFR or Toll-like receptor family members. TRAF3 is reported to be a highly versatile regulator that positively controls type I interferon production, but negatively regulates MAP kinase activation and alternative nuclear factor-κB signalling [Cai et al., 2013]. TNIK, in contrast, is a member of the germinal center kinase family, and is widely involved in cytoskeleton organization, neuronal dendrite extension, cell proliferation, and glutamate receptor regulation. Notably, an emerging body of evidence suggests that TNIK is a novel activator of Wnt signalling [Yu et al., 2014], and the Wnt pathway plays critical roles in the regulation of many cellular functions, including cellular proliferation, differentiation, migration, and maintenance of pluripotency. However, although GWAS linked variants have been reported with respect to TNIK [Zagajewska et al., 2018]; to our knowledge, changes in regulation of TNIK protein has not been reported in any previous glaucoma study.

In this report, we have shown that high IOP subjected rat retinas demonstrated shared components of several tissue stresses, and adaptive or neuroprotective responses that were also identifiable in human glaucoma subjects [Mirzaei et al., 2017a]. Notably, many biochemical networks such as modulation of cholesterol transport proteins and classical complement activation were uniquely affected in the human eyes [Mirzaei et al., 2020]. It is important to emphasise that the responses detected in rat retinas were caused by IOP increase alone without any underlying manifest neurodegenerative or vascular pathology. This is a limitation of the current glaucoma models being used in research, nevertheless, the molecular data helps to dissect various components of the glaucoma pathology as a complex neurodegenerative/ vascular disorder. In this study we have analysed the proteome of the whole retinal tissues, which limits our ability to distinguish the individual contribution of distinct cell types that comprise the retina. This limitation has been extensively discussed in our previous study analysing the human glaucoma retinal tissues [Mirzaei et al., 2017a]. Further, a different experimental design in which animal retinal tissues are harvested at intermediate time points will help us understand the chronology of biochemical changes that occur in response to chronically elevated IOP exposure.

In conclusion, the proteomics data provides new insights into a purely high IOP induced scenario for retinal changes, without the complex interplay of ageing, genetic or systemic factors and drug effects, which are inevitable confounding variables in human cohort studies. Future research will help elucidate the individual effects of various risk factors in glaucoma, to help us better understand the disease process and thereby develop new therapeutic strategies.

## 4 Material and Methods

### 4.1 Animals

Adult male Sprague-Dawley rats (n=10) were used for the experiments (High IOP/glaucoma: 5, control: 5). All animals were maintained at controlled temperature (21 ± 2°C) for 12-hour light/dark cycles. All procedures were performed in accordance with the Australian Code of Practice for the Care and Use of Animals for Scientific Purposes and the guidelines of the ARVO Statement for the Use of Animals in Ophthalmic and Vision Research. The animals were subjected to anaesthesia using intraperitoneal (i.p.) administration of ketamine (75 mg/kg) and medetomidine (0.5 mg/kg)[You et al., 2012] for the microbead injections[Chitranshi et al., 2018b].

### 4.2 Intra-ocular injections and pressure measurements

An experimental glaucoma model was established by producing a chronic increase of intra-ocular pressure (IOP) in mice by microbead (Fluospheres, Molecular Probes, 10 μm) injections in the anterior chamber as reported previously [Chitranshi et al., 2019; Gupta et al., 2014; You et al., 2014]. Intraocular injections (3.6× 10^6^ microbeads/mL) in the anterior chamber of the eye were performed until a sustained increase in IOP was observed. Eyes were injected using a 25-μL Hamilton syringe connected to a disposable 33-gauge needle (TSK, Japan). All intracameral procedures were performed under magnification using an operating microscope (Carl Zeiss, Germany) with care taken to avoid needle contact with the iris or lens. The needle was inserted bevel down, tangentially beneath the corneal surface, to facilitate self-sealing of the puncture wound. Once the needle tip was visualised within the anterior chamber, 2 μL microbead solution was injected. At the end of the procedure, anaesthesia was reversed using atipamazole (0.75 mg/kg subcutaneous injection, and 0.3% ciprofloxacin drops (Ciloxan; Alcon, Australia) and 0.1% dexamethasone eye drops (Maxidex, Alcon) were instilled in both eyes. An ointment (Lacri-lube; Allergan, Australia) was also applied to protect against corneal drying until the animal recovered. During anaesthesia and prior to each injection, IOP was measured by using a handheld electronic tonometer (Icare Tonovet, Finland). The IOP displayed on the tonometer was the mean of six consecutive measurements. Three consecutive IOP readings were obtained from each eye and the average number was taken as the IOP for that time point.

### 4.3 Preparation of protein samples

The retinas of experimental glaucoma (n=5) and control animals (n=5) were extracted from the eyes using the surgical microscope. Protein extraction was carried out using lysis buffer (20 mM HEPES, pH 7.4, 1% Triton X-100, 1 mM EDTA) containing protease and phosphatase inhibitors. Samples were subjected to probe sonication (3 pulses/15s with 20s between each pulse). Lysed samples were centrifuged (15,000 g for 10 min at 4°C) and protein in the supernatant concentrated by precipitation using a chloroform methanol precipitation procedure [Wessel and Flügge, 1984]. Precipitated protein was resuspended in 200 ul of 8M Urea in 50 mM Tris (pH 8.8). Protein concentration was determined using a BCA assay (Pierce, Rockford, USA) and bovine serum albumin (BSA) as a standard. Solubilised proteins were reduced using 5mM DTT and alkylated using 10mM iodoacetamide. Proteins (150 μg) were initially digested at room temperature overnight using a 1:100 enzyme-to-protein ration using Lys-C (Wako, Japan), followed by digestion with Trypsin (Promega, Madison, WI) for at least 4 hours at 37 °C also at a 1:100 enzyme-to-protein ratio.. Resultant peptides were acidified with 1% TFA and purified using SDB-RPS (Empore) Stage Tips [Kulak et al., 2014].

### 4.4 TMT labelling and liquid chromatography electrospray ionization tandem mass spectrometry (LC-ESI-MS/MS)

Details of the 10 plex TMT experimental workflow are shown in figure 1. Briefly, peptides (70 μg) from each retinal sample were subjected to TMT labelling as outlined in our previous study on human glaucoma retinal tissues [Mirzaei et al., 2017a]. Briefly, pooled and labelled peptides were separated into 12 fractions following HpH chromatography and dried by vaccum centrifugation. Peptide fractions were analysed by nano-LC-MS/MS using an ultra-high-pressure liquid chromatography system (Proxeon) and an Orbitrap Fusion Tribrid-MS (Thermo Scientific, USA) mass spectrometer. Peptide fragmentation was performed using the (SPS)-MS^3^ method described previously [McAlister et al., 2014].

### 4.5 Database searching/quantification and statistical analysis

Charge state and monoisotopic m/z values were corrected using a method previously detailed by Huttlin et al [Huttlin et al., 2015]. Spectra were search against an indexed *Rattus norvegicus* Uniprot database using the Sequest search algorithm and included carbamidomethylation of cysteine residues, and TMT labelling of peptide N-termini and lysine residues as static modifications, oxidation of methionine as a dynamic modification, and a precursor ion tolerance of 20 ppm and fragment ion tolerance of 0.8 Da. A linear discriminant analysis was used to filter Sequest matches to a false discovery rate (FDR) of 1% at the peptide level based on matches to reversed sequences [Elias and Gygi, 2010]. Protein were ranked by multiplying peptide probabilities and the dataset was finally filtered to 1% protein FDR. TMT reporter ion quantification was assessed using the strategy described previously [Chick et al., 2016; McAlister et al., 2012]. Peptides with a total signal-to-noise (S/N) for all channels of >200 with a precursor isolation specificity of >0.75 were used for quantification. TMT reporter ion intensity values were normalized by summing values across all peptides within each channel then correcting each channel to the same summed value. Protein level quantification was performed using normalized S/N values for all peptides assigned to a given protein. Differentially expressed proteins were assessed using a two-sample t-test (p ≤0.05) and a fold change threshold (≥ 1.2 for up-regulation or ≤ 0.83 for downregulation). Reproducibility of the TMT experiment was evaluated further using our in-house ‘TMTPrepPro’ software reported previously [Mirzaei et al., 2017b].

### 4.6 Bioinformatics and functional pathway analysis

Functional pathway enrichment and protein-protein interaction analysis of differentially modulated proteins were performed using Ingenuity pathway analysis (IPA) and STRING protein-protein network analysis [Szklarczyk et al., 2015] as described previously [Mirzaei et al., 2017a]. For Ingenuity, the canonical pathway analysis highlighted the function-specific genes present in each networks. STRING Cytoscape plug-in assisted in detecting the protein-protein interactions as well as classifying the differentially modulated pathways and networks.

### 4.7 Tissue Sectioning and Immunofluorescence Analysis

Rats eyes were enucleated following the transcardial perfusion with 4% paraformaldehyde. The eyeballs were further subjected to PFA treatment for 1 h. The eye tissues were rinsed 3 times with 0.9 % saline and dropped into 30% sucrose solution till the tissues sank to the bottom. The eyeballs were embedded in OCT media and 10-15-μm thick retinal sections were cut with the help of cryostat and dipped into ice-cold ethanol for tissue permeabilization for antibody treatment. The sections were blocked with serum treatment and subjected to incubation with primary antibodies (1:300). The eye sections were subsequently incubated with secondary antibodies (1:400 in Tris phosphate saline buffer) as indicated for 1 hour in the dark room and mounted using antifade media containing DAPI for nuclear staining. The images were obtained using Carl Zeiss microscopes and data analysed. PFA fixed human eyes were similarly washed with 0.9% saline and embedded in OCT media. The sections were cut using cryostat, stained with crystalline antibodies and images captured as described above.

## 5 Conclusion

In present study, we aimed to investigate shared retinal protein components from experimental glaucoma and human glaucoma subjects. We identified that experimentally elevated IOP in rat retinas shared components of several tissue stresses and adaptive or neuroprotective responses that were also identifiable in post-mortem retinal tissue from human glaucoma subjects. Notably, many biochemical networks such as modulation of cholesterol transport proteins and classical complement activation were uniquely affected in the human eyes. It is important to emphasise that the responses detected in rat retinas were caused by IOP increase alone without any underlying manifest neurodegenerative or vascular pathology. This is a limitation of the current glaucoma models being used in research; nevertheless, the molecular data helps to dissect various components of the glaucoma pathology as a complex neurodegenerative/ vascular disorder. In addition, the proteomics data provides new insights into a purely high IOP induced scenario for retinal changes, without the complex interplay of aging, genetic or systemic factors and drug effects, which as inevitable confounding variables in human cohort studies. Future research will elucidate the individual effects of various risk factors in glaucoma and help us to better understand the disease process and thereby develop new therapeutic strategies.

## Acknowledgments

We acknowledge funding support from the Australian Government National Collaborative Research Infrastructure Scheme (NCRIS), National Health and Medical Research Council (NHMRC) Australia, Ophthalmic Research Institute of Australia (ORIA) and Macquarie University, NSW, Australia.

